# Exploring the Role of PEDOT Electrodes in Electrostimulation of Vascular Endothelial Cells

**DOI:** 10.1101/2025.01.30.635422

**Authors:** Jan Víteček, Matouš Kratochvíl, Martin Weiter

## Abstract

The potential of electrical stimulation for cell control in tissue engineering remains largely underestimated, as does the use of organic semiconductors. The tunable physico-chemical properties of organic semiconductors offer advantages over commonly used inert metals.

This study investigates the effects of pulsed electrostimulation and electrode materials, gold and poly(3,4-ethylenedioxythiophene) (PEDOT), on the physiological functionality of human vascular endothelial cells. A novel electrostimulation platform incorporating gold or PEDOT electrodes was developed and characterized for electrical performance. Human umbilical vein endothelial cells were cultured on this platform, and the effects of electrostimulation and electrode material were assessed through morphological analysis, nitric oxide (NO) production, and expression of key endothelial marker genes.

PEDOT electrodes produced higher electrical current during electrostimulation. Interestingly, cell morphology, including elongation and alignment, showed minimal changes under electrostimulation. NO production, a key marker of vascular health, was enhanced by electrostimulation, with PEDOT electrodes showing a trend toward greater NO accumulation than gold. Gene expression analysis revealed material- and stimulation-specific trends: electrostimulation generally upregulated KLF2, KLF4, and CYP1B1 on PEDOT electrodes but suppressed their expression on gold electrodes.

These findings suggest that PEDOT electrodes, with their enhanced electrochemical properties and ability to support endothelial functionality, provide a safe and efficient platform for endothelial cell electrostimulation. This study advances understanding of the interplay between material properties and electrostimulation and highlights PEDOT as a promising candidate for vascular tissue engineering and regenerative medicine.

## Introduction

Cells interact with their environment through a variety of mechanisms, including electrically mediated processes. These complex interactions play a critical role in controlling cell behavior (1–4). However, the potential of electrical stimulation for cell control in tissue engineering remains largely underestimated, as does the use of organic semiconductors in this field (5). This gap is partly due to the lack of materials capable of optimally supporting interactions between organic semiconductors and cells. Despite significant progress, the utilization and performance of organic conductive materials in cellular applications remain relatively limited. One of the primary challenges is the incomplete understanding of the cellular and molecular mechanisms governing the interactions between these materials and biological systems (6,7).

Our research focuses on vascular physiology, where vascular endothelial cells play a pivotal role. These cells form the luminal layer of blood vessels, constituting the primary barrier between the bloodstream and surrounding tissues. This layer is critically important for cardiovascular function, as it regulates blood pressure, clotting, nutrient and water transport, and immune responses. Dysfunction of the endothelium can lead to severe conditions, including atherosclerosis, myocardial infarction, and stroke. Notably, endothelial function is maintained through its ability to perceive blood flow, which drives cell proliferation, migration, and spatial organization (8,9). A fully functional vascular endothelium is characterized by nitric oxide (NO) production, a key mediator of vasodilation essential for maintaining vascular tone (10). Additionally, the endothelium exhibits specific mechanisms of stress protection in which cytochromes appear to be involved (11). The molecular basis of endothelial function is regulated by transcription factors such as KLF2 and KLF4, which modulate the expression of genes critical for proper vascular activity (12).

Relatively few studies have examined the effects of electrostimulation on endothelial cells. Early work demonstrated that electrical stimulation can influence endothelial morphology, migration, and orientation within an electric field (4). Furthermore, the potential role of electrostimulation in promoting endothelial repair during tissue healing has been proposed (13). However, the precise effects of electrostimulation on endothelial cell physiology and the molecular mechanisms involved remain poorly understood.

To conduct cell electrostimulation most previous studies have employed metal electrodes, often at voltages exceeding the electrolytic limit of the extracellular medium, necessitating the use of agar bridges to avoid damage (14). Organic semiconductors offer a promising alternative, as they enable efficient electrostimulation below the electrolytic limit, allowing for direct contact with cells. These materials may also provide additional, material-specific cues that influence cell behavior.

In the present study, we developed a novel platform for the electrostimulation of endothelial cells using inert metal electrodes (gold) and organic semiconductor electrodes (PEDOT). The functionality of these electrodes was carefully characterized. This platform was then applied to investigate whether the combination of electrostimulation and electrode material could enhance endothelial cell functionality.

## Materials and methods

### Electrostimulation platform

The platform was prepared using the layout of a standard microscope slide. The electrodes 8 x 40 mm were centered to the slide with 1 mm spacing between them. First the gold electrodes were rendered by means of physical vapour deposition (15) This platform was further referred to as the gold platform.

For the electrochemical polymerization of PEDOT films on gold support a galvanostatic method was adopted. The galvanostatic polymerization was carried out in 2 electrode arrangement with Pt coil as a counter electrode and thin film of gold thermally evaporated on a glass slide as working electrode. The used current density was 0.2 mA·cm^−2^ (with current value given by the area of electrode) and deposition time was 300 s. The electrolyte consisted of the EDOT monomer (5·mmol l^−1^) and tetrabutylammonium hexafluorophosphate (TBAPF6, 100 mmol l^−1^) dissolved in anhydrous acetonitrile. The electrolyte was deoxygenated by bubbling argon into the cell. This platform was further referred to as the PEDOT platform. The platform was completed with a plastic grid forming two slots with 2 cm^2^ culture area and one slot for wiring connection.

### Platform characterization

In addition to optical microscopy in reflected light mode the standard scanning electron microscopy was carried out to explore the surface features of electrodes (described elsewhere, e.g (16))

The capacitive properties of PEDOT coated electrodes were studied by cyclic voltammetry (CV). The measurement was conducted using a pocketSTAT2 potentiostat (Ivium Technologies B.V.) in True Linear Scan mode, thus applying analogue potential. The set-up for cyclic voltammetry was a three-electrode electrochemical system, consisting of a golden film on glass slide coated with PEDOT as a working electrode, an Ag/AgCl reference electrode (Metrohm AG) and a Pt coil counter electrode. The electrolyte used was a phosphate-buffered saline (PBS). Prior to each measurement the electrolyte was deoxygenated by bubbling nitrogen into the measurement cell. Cyclic voltammograms were recorded by scanning potential from −200 to 200 mV vs Ag/AgCl RE for studying the capacitive properties or from 0 to −1.5 V vs Ag/AgCl RE in case of essential characterization. The scanning rate was 50 mV·s^-1^ in all measurements.

The platform wiring connections as well as the capacitance of electrode systems were routinely checked by means of MT-1710 multimeter (Pro’s Kit, CZ).

### Electrostimulation pulses

Alternating rectangular wave pulses were applied. One cycle lasted one second as follows 0.2 V for 25 ms, -0.2 V for 25 ms, delay 950 ms. The voltage pulses were achieved by means of a pocketSTAT2 potentiostat/galvanostat (Ivium Technologies, Eindhoven, NL) in two electrode operation mode.

### Hydrogen peroxide assay

One hundred microliters of sample was mixed with 80 µl of 2.5 mmol l^-1^ M 2,2’-azino-bis(3-ethylbenzothiazoline-6-sulfonic acid (ABTS) in physiological buffered saline (pH 7.4) and 20 µl 0.1 mg ml^-1^ horse radish peroxidase. The absorbance was read using a spectrophotometer at 414 nm after 10 minutes. Such approach has quantification limit about 1 µmol l^-1^ hydrogen peroxide.

### Cell culture

The platform was coated with collagen IV as indicated in (16). Human umbilical vein endothelial cells (C-12253, Merck, DE) were cultured in EGM-2 medium as previously (17). Two days prior to electrostimulation each chamber in the electrostimualtion platform was loaded with 80 to 100 thousands of cells. The culture medium was exchanged two hours before electrostimulation. The electrostimulation was carried out as above.

### Image analysis of cell orientation and elongation

The determination of cell orientation and elongation was carried out by means of image analysis (16).

### Nitrite accumulation

The production of nitrite was estimated by means of Griess reagent (G4410, Merck) according to manufacturer instruction. The quantification limit of the approach was about 0.5 µmol l^-1^.

### Gene expression

Total RNA was extracted using the CatchGene Tissue DNA Kit (CatchGene; Taipei, Taiwan). Complementary DNA was synthesized according to the manufacturer’s instructions for the gb Elite Reverse Transcription kit (Generi Biotech) using TurboCycler Lite thermal cycler (BlueRay Biotech, Taipei, Taiwan). Real-time quantitative PCR (RT-qPCR) reactions were performed in a LightCycler480 instrument using LightCycler480 Probe Master solutions (Roche; Basel, Switzerland). The following program was used: initial denaturation step at 95 °C for 10 min, followed by 45 cycles (95 °C for 10 s, 60 °C for 30 s, and 72 °C for 1 s) and the final cooling step at 40 °C for 1 min. β-actin (ACTB) gene was used as reference gene for both synthetic primers and taqman probes. The expression of genes of interest was presented as 2^−Δcq^. Synthetic primers for genes KLF2, KLF4, ASS1 (Tab. 1) were designed using the Universal Probe Library Assay Design Center together with appropriate probe. Taqman probes (see Assay ID in Tab. 1) of CYP1B1 gene were obtained from Thermofisher Scientific.

**Table 1.**
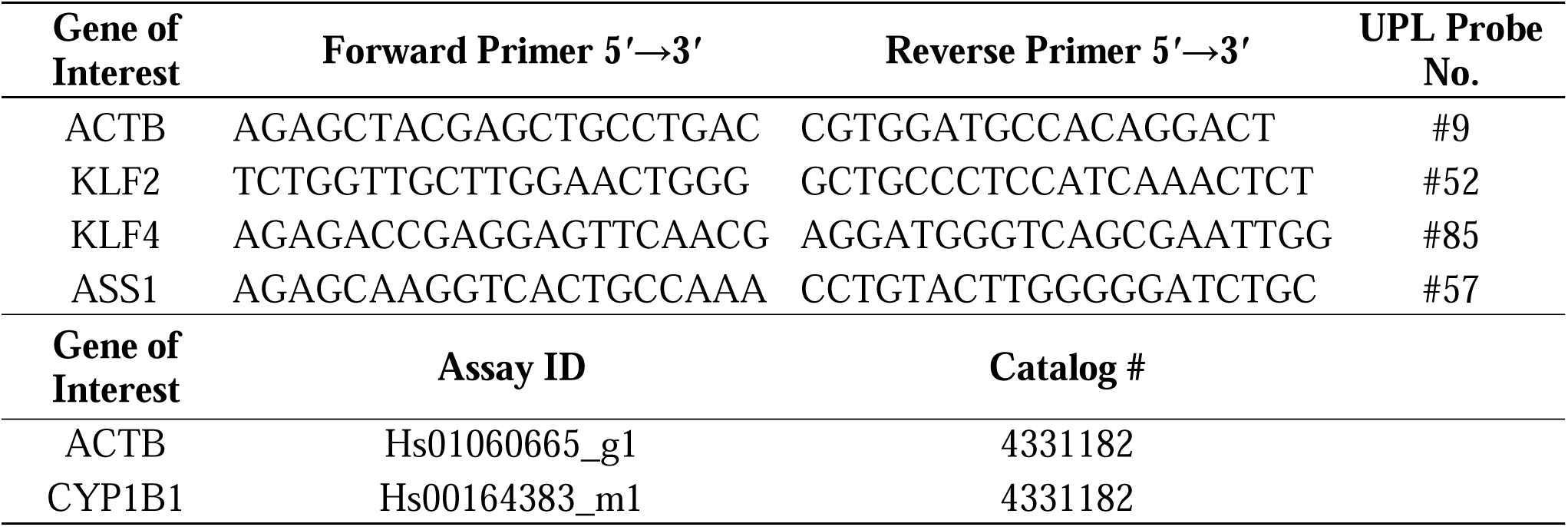
Primers, probes and reagents used for the real-time quantitative PCR.

### Data processing and statistics

Data were processed in Microsoft Excel (2016) and are expressed as mean ± standard error of mean (SEM). The statistical analysis was performed in STATISTICA 12 (StatSoft, Dell Inc., USA). Unpaired t-test and one-way ANOVA with post-hoc Fischer’s LSD test were used to compare the data. P-values ≤ 0.05 were considered to be statistically significant.

## Results

### Electrostimulation platform and its characterization

The electrostimulation platform was designed based on layout of standard microscope slide to ensure modularity. The electrodes were made by means of physical vapor deposited gold (further referred as “gold electrode”) or the gold served as a support for electrochemical polymerization of EDOT to deposit PEDOT (further referred as PEDOT electrode). (Fig. 1A).

**Fig. 1.**
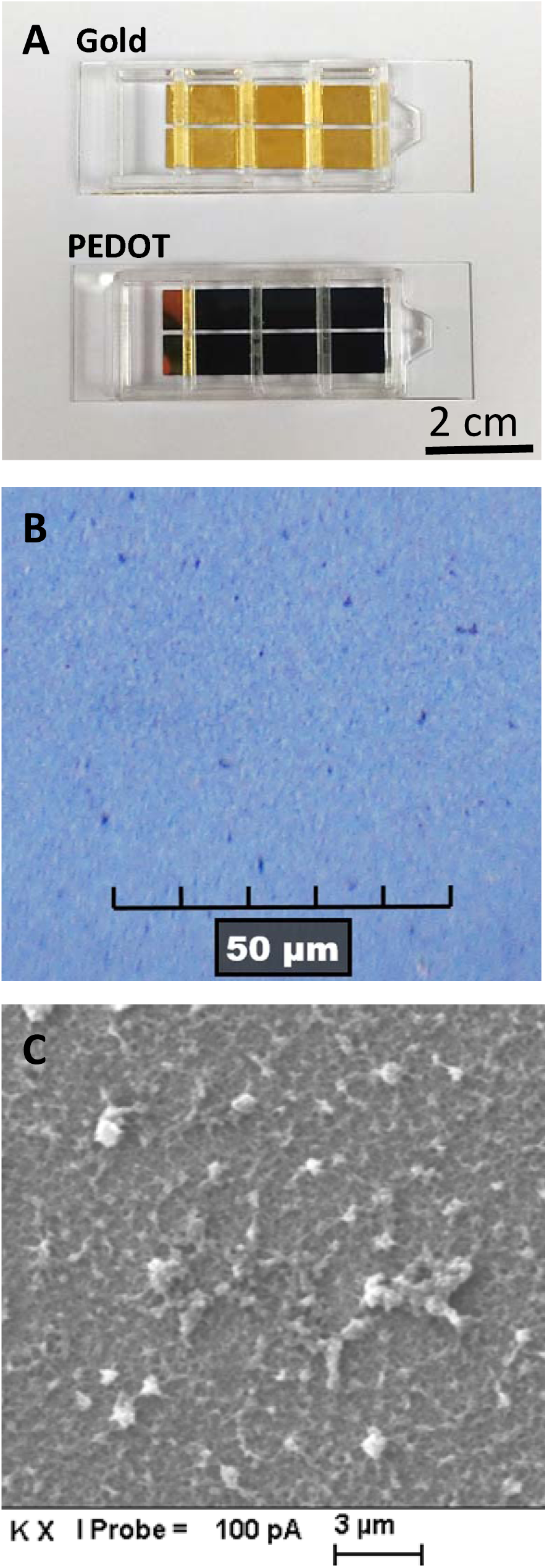
Electrostimulation platform and its microscopic characterization: The platform was designed on a layout of standard microscope slide (26 per 76 mm). Two paralel electrodes were created by physical vapour deposition of gold with spacing of 1 mm – gold electrodes. For PEDOT electrodes platform the material was deposited by electropolymerization. Plastic grid was used to provide cell culture space and cut-out for wiring the electrode system (**A**). Reflected light microphoto of PEDOT electrode surface (**B**). Scanning electron microscopy of the PEDOT electrode surface (**C**). Microscopic images show representative data of three independent PEDOT electrode preparations.

The surface of electrodes was characterized. The gold electrode was without major roughness seen in optical and electron microscopy (not shown). Whereas the surface of PEDOT electrode prepared by monomer (EDOT) electropolymerization showed somewhat granular structure in optical microscopy (Fig. 2B). The electron microscopy supported such an observation as it revealed substantial roughness at submicrometer scale. Further certain degree of porosity at the level of 100 nm can be inferred from the image (Fig. 2C).

**Fig. 2.**
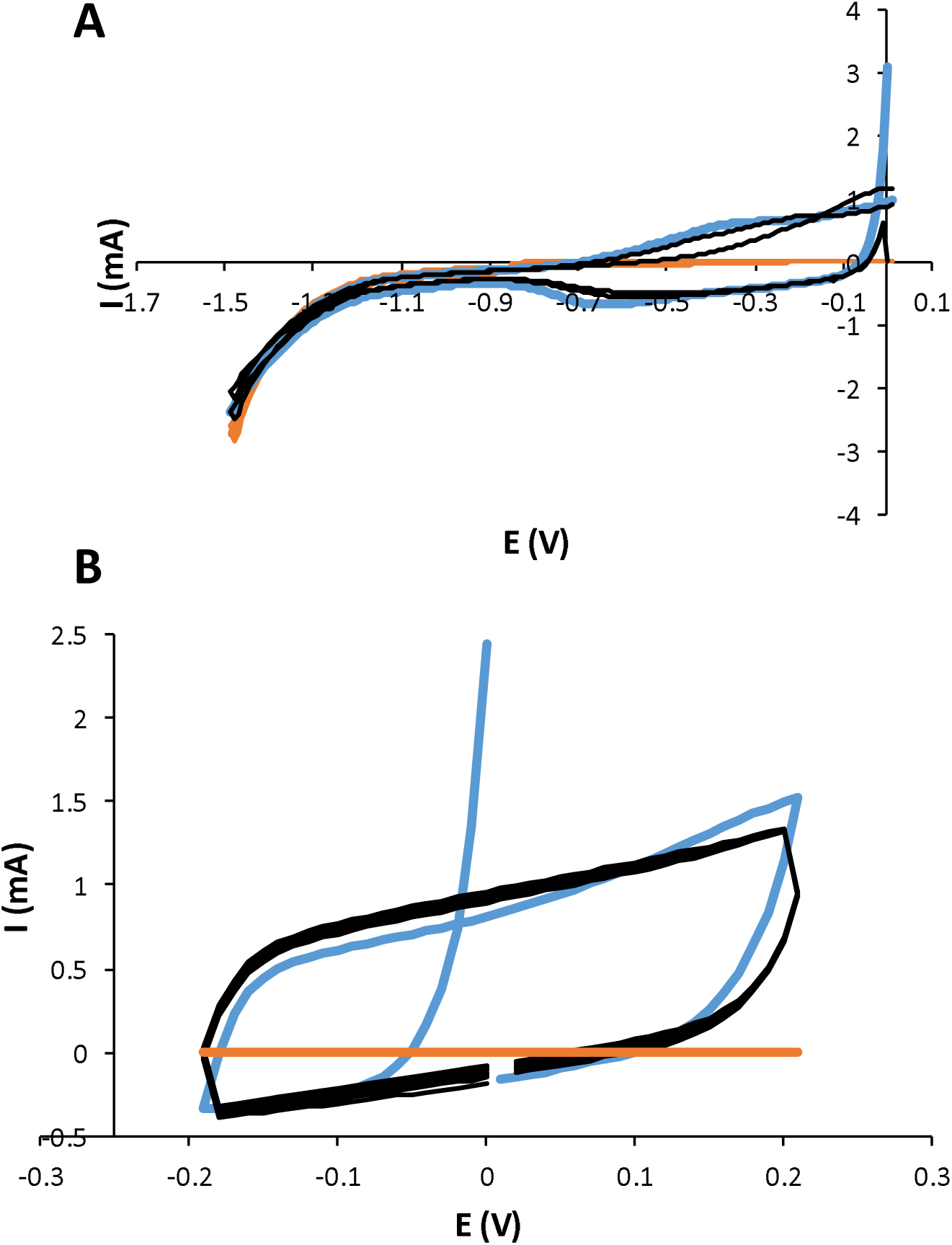
Cyclic voltammetry characterization of the gold and PEDOT electrodes: Working electrodes with area 2 cm^2^ at scan speed 50 mV/s were used. Three cycles of PEDOT and gold electrode analysis in the range from 0 to -1.5 V (**A**). Five cycles of of PEDOT and gold electrode analysis in the range from -0.2 to + 0.2 V (**B**). **Blue line** first cycle of PEDOT, **black lines** remaining cycles for PEDOT. Orange lines – cycles of gold.

To provide thorough characterization of the electrode systems, first the voltammetric characterization was carried out within the range of 0 to -1.5V to provide characteristics over broader range of voltages. The gold electrode produced very low currents up to about -0.7 V, where the current started to increase due to onset of physiological medium electrolysis. The PEDOT electrode produced, however, more pronounced currents right from the beginning. At about -0.7 V the absolute value of the current decreased and gradually reached a similar level at -0.9 V like in the case of gold electrode. Further a change in course of follow up cycles indicated electrochemical alteration of the PEDOT electrode material (Fig. 2A).

To verify suitability of PEDOT electrodes for the intended electrostimulation mode (see above) the cyclic voltammetry characterization was carried within the range of -0.2 to + 0.2 V. After the first cycle we have obtained symmetrical voltammograms indicating dominant contribution of capacitive effect of the PEDOT electrodes and possible reversible electrochemical events. On the other hand, gold electrodes showed negligible electrical currents across whole cycle (Fig. 2B).

The capacitance of electrode systems in the platform was 10±1 µF for gold 1.4±0.2 mF for PEDOT vs. 10±1 µF for gold.

### Electrostimulation mode

The electrostimulation mode intended for cells derived from our previous experience (15): 0.2 V, 25 ms; -0.2 V, 25 ms, delay 950 ms was thoroughly checked for the current response of the electrostimulation system. Thus, the peak current of gold electrodes ranged up to 0.6 mA. Further, the electrode system charged very quickly as indicated by immediate dropdown of current after change of electrical potential. In case of PEDOT electrodes the peak current was much more pronounced – up to about 2 mA. The charging of the electrode system was much slower as the current decreased gradually within the polarisation phase and did not reach equilibrium. (Fig. 3A, B). Since the electrochemical processes at the surface of electrodes may result in production of active oxygen species in aerobic environment even at low voltages (18) we meticulously checked this parameter. The representative form of active oxygen species – hydrogen peroxide – way assayed within physiological medium after up to 2 hours since start of electrostimulation. The approach with quantification lever of 1 µM hydrogen peroxide provided negative results for both gold and PEDOT electrodes.

**Fig. 3.**
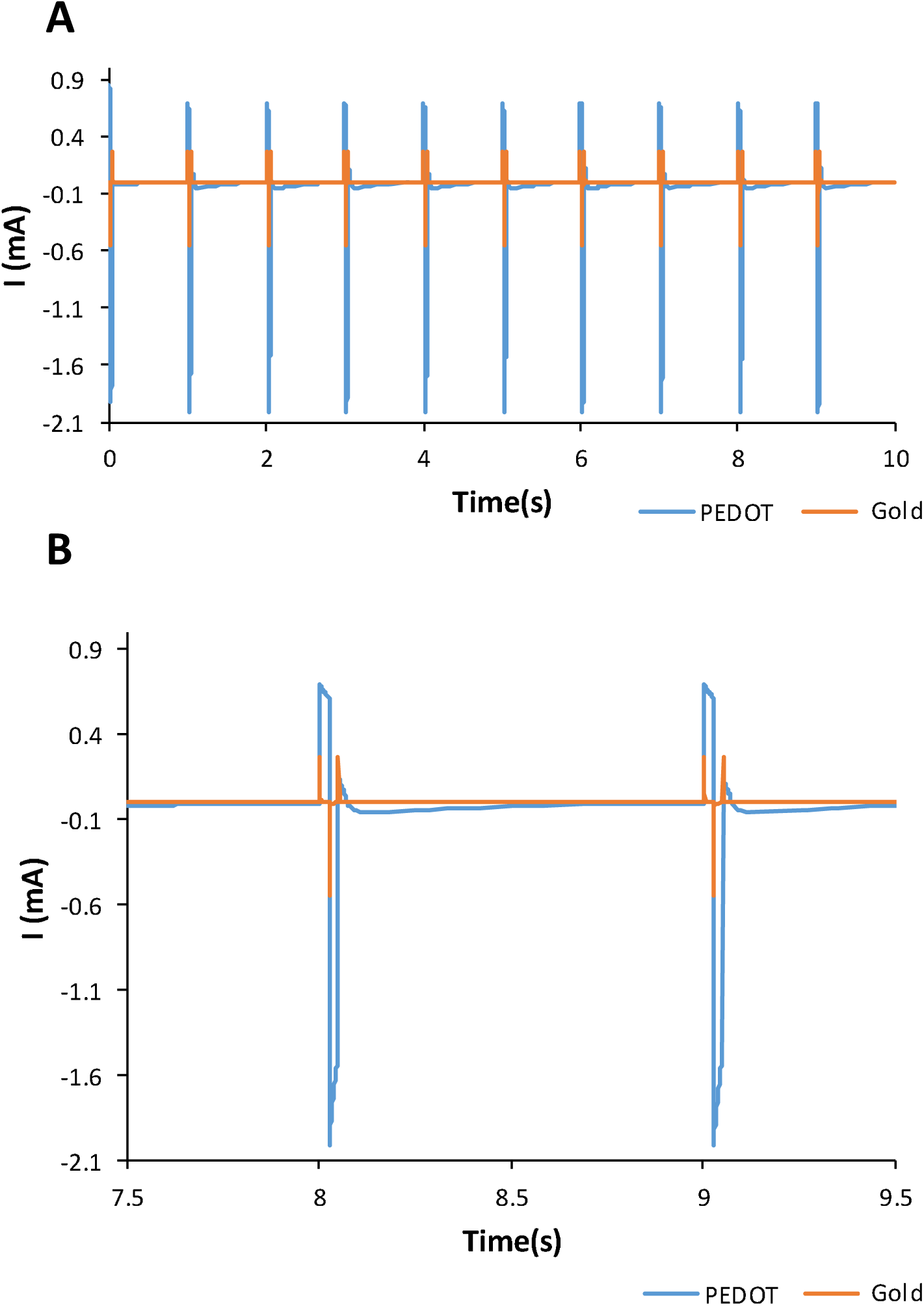
Time course of current pulsation during electrostimulation. The voltage pulses (0.2 V, 25 ms; -0.2 V, 25 ms, delay 950 ms) were applied as in electrostimulation of cells. Typical time course is displayed (**A**). A magnified cut-out showing more details is provided (**B**).

### Cell electrostimulation

Before the electrostimulation of cells, the platform biocompatibility was verified meticulously. The platform supported cell growth attatchment and spreading to the same level as standard cell culture plastics (data not shown) if coated with collagen IV. The cells were electrostimulated for 48 hours. The electrostimulation did not affect cell viability as determined by double staining with fluorescein diacetate and propidium iodide what corresponds with our previous experience (15).

### Cellular morphology

In order to determine the impact of electrostimulation and electrode material on morphology of HUVEC, image analysis of cell orientation and elongation was carried out. Cells mostly showed no preferential orientation in regard to electrodes, though there was a marginal trend when electrostimulated with PEDOT electrodes. However, the PEDOT electrode stimulated cells showed no clear orientation along with electrodes (Fig. 4). Similarly to the above, the elongation of cells expressed as an elongation index did not show any clear-cut effect of electrostimulation as well as the electrode material (Fig. 5)

**Fig. 4.**
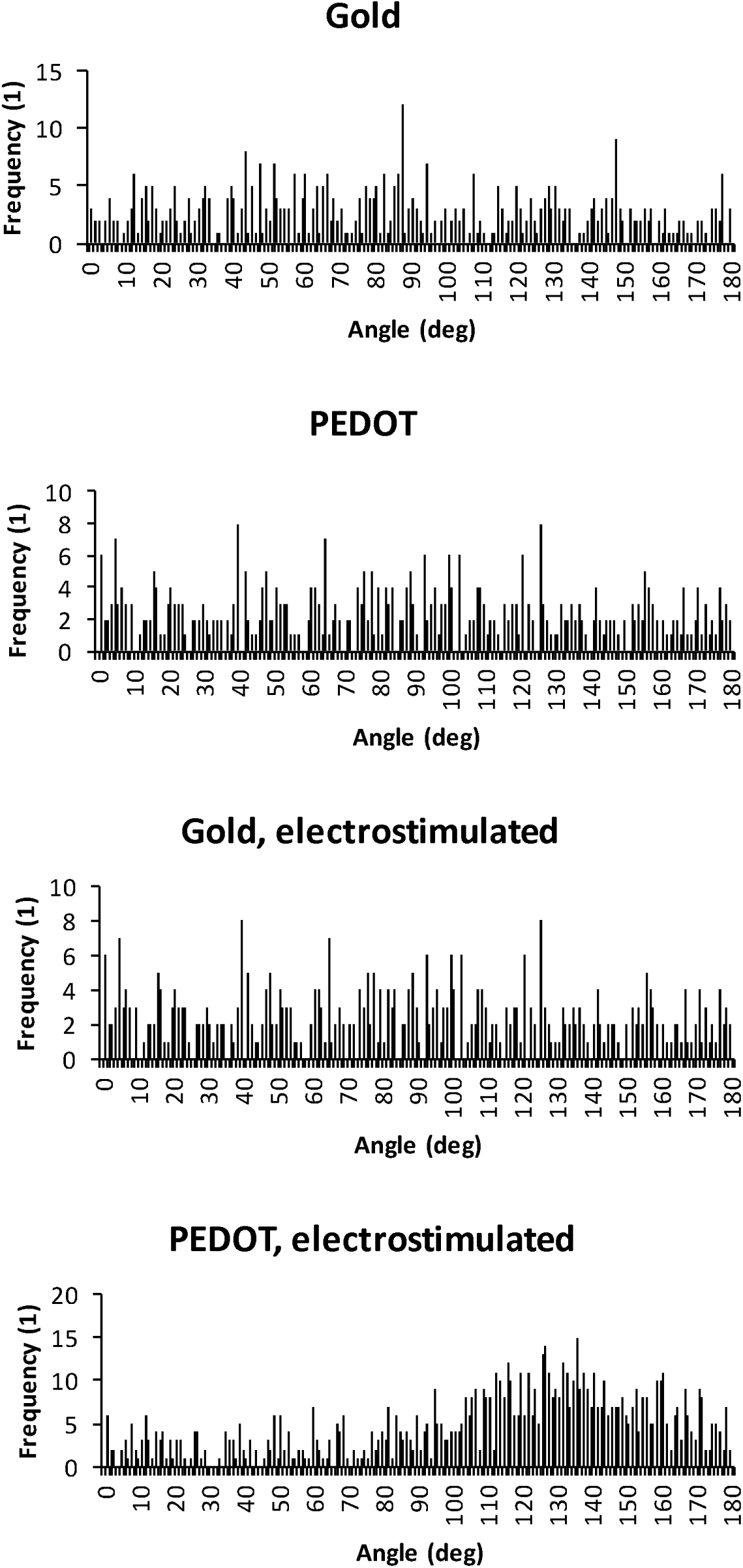
Analysis of cell orientation. Human endothelial cells (HUVEC) were electostimulated for two days. The cell orientation was assayed in microscopic images by image analysis (ImageJ) as an orientation of major axis of an ellipse circumscribed to each cell. Each histogram provides data for at least 200 cells. Representative data out of three biological replicates are shown.

**Fig. 5.**
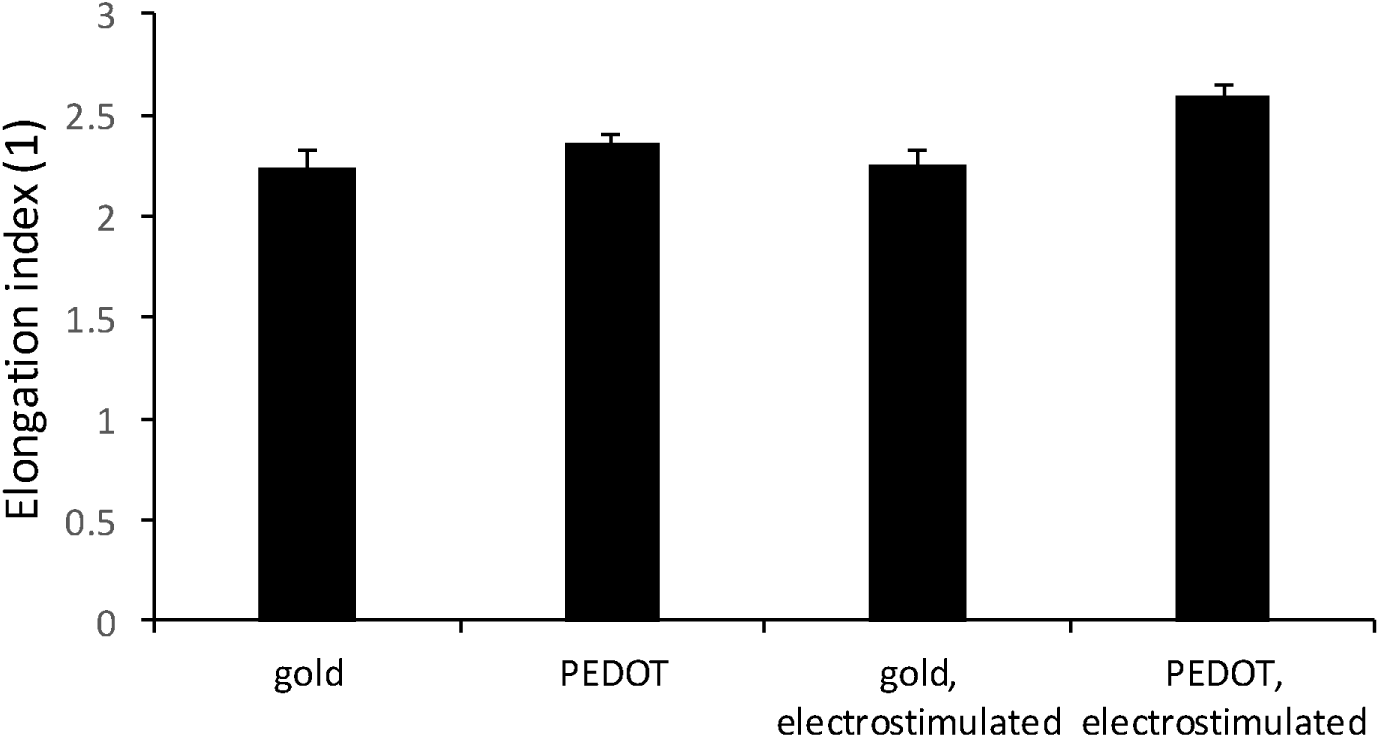
Analysis of cell elongation. Human endothelial cells (HUVEC) were electostimulated for two days. The cell elongation was assayed in microscopic images by image analysis (ImageJ) as aspect ratio of an ellipse circumscribed to each cell. Each data point shows mean value ± SEM for at least 200 cells. Representative data out of three biological replicates are shown.

### Nitric oxide production

Nitric oxide (NO) production is one of vital signatures of endothelial cells. As a readout the accumulation of nitrite was assayed. Since the assay was conducted very close to quantifiable concentrations a standard addition approach was carried out to exclude the interference of conditioned media from different experimental variants. The electrostimulation with gold as well as PEDOT electrodes showed clear accumulation of nitrite. There was, however, no clear-cut difference between electrostimulation with gold and PEDOT electrodes though PEDOT ones indicated a trend towards improvement of nitrite accumulation (Fig. 6).

**Fig 6.**
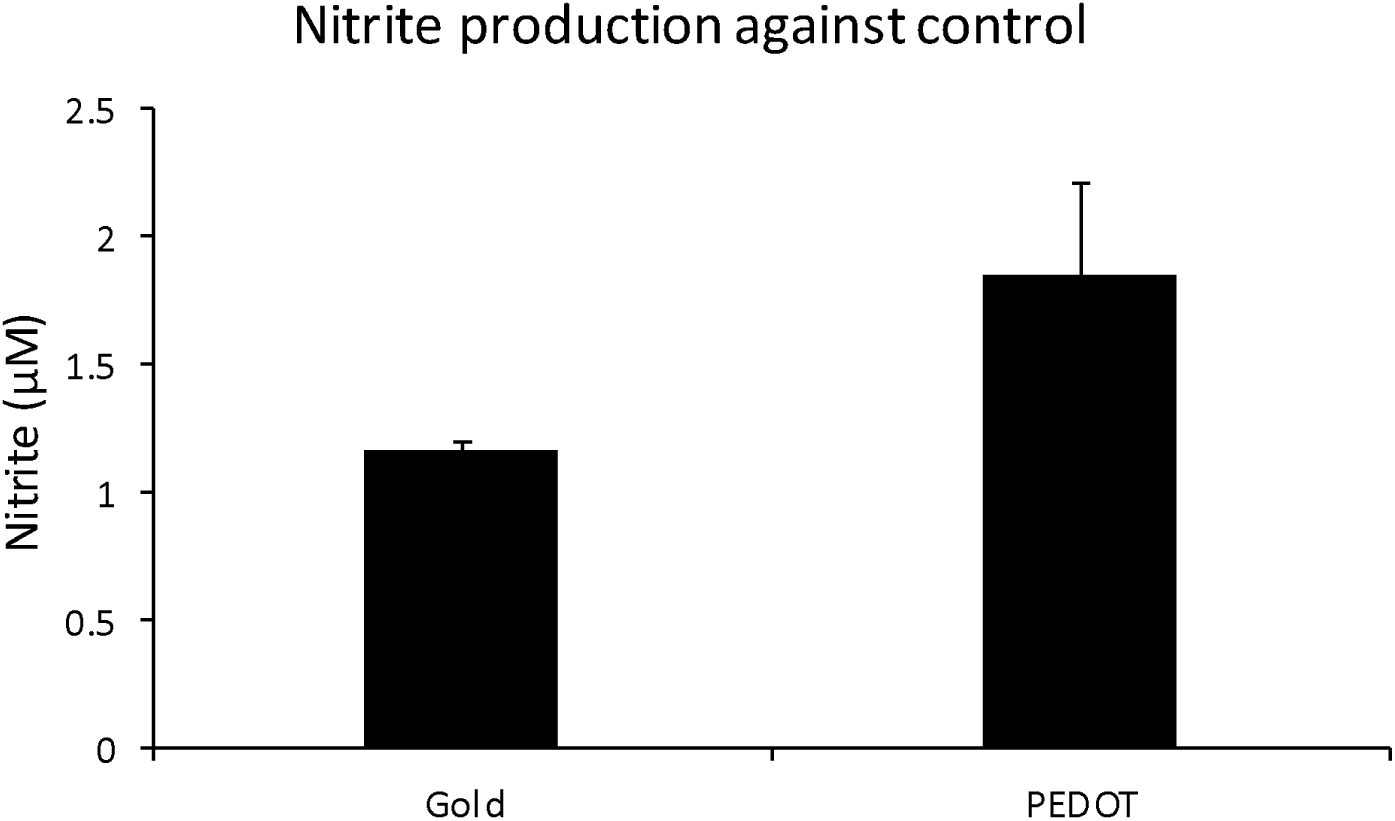
Determination of nitrite after electrostimulation. Human endothelial cells (HUVEC) were electostimulated for two days. Nitrite was determined by Griess reagent. Differences between appropriate electrostimulated and control variants are displayed. The control variants were at level of about 2 µM. Data are indicated as mean value ± SEM, N=3. There was no significant difference (p 0.12).

### Expression Analysis of Endothelial Marker Genes

To explore key aspects of endothelial cell function, we assessed the expression of selected marker genes at the mRNA level. These included KLF2 and KLF4 (general markers of endothelial function(12)), ASS1 (involved in NO metabolism (12)), and CYP1B1 (indicative of stress protection (11)). While the results showed observable trends, differences were not statistically significant.

To draw a good comparison, the effect of electrostimulation using PEDOT electrodes versus gold electrodes was first evaluated against appropriate controls. Subsequently, the effect of PEDOT electrodes versus gold electrodes, with and without electrostimulation, was analyzed to provide more detailed insights into the impact of the material itself.

#### KLF2 Expression (Fig. 7A)

**Fig 7.**
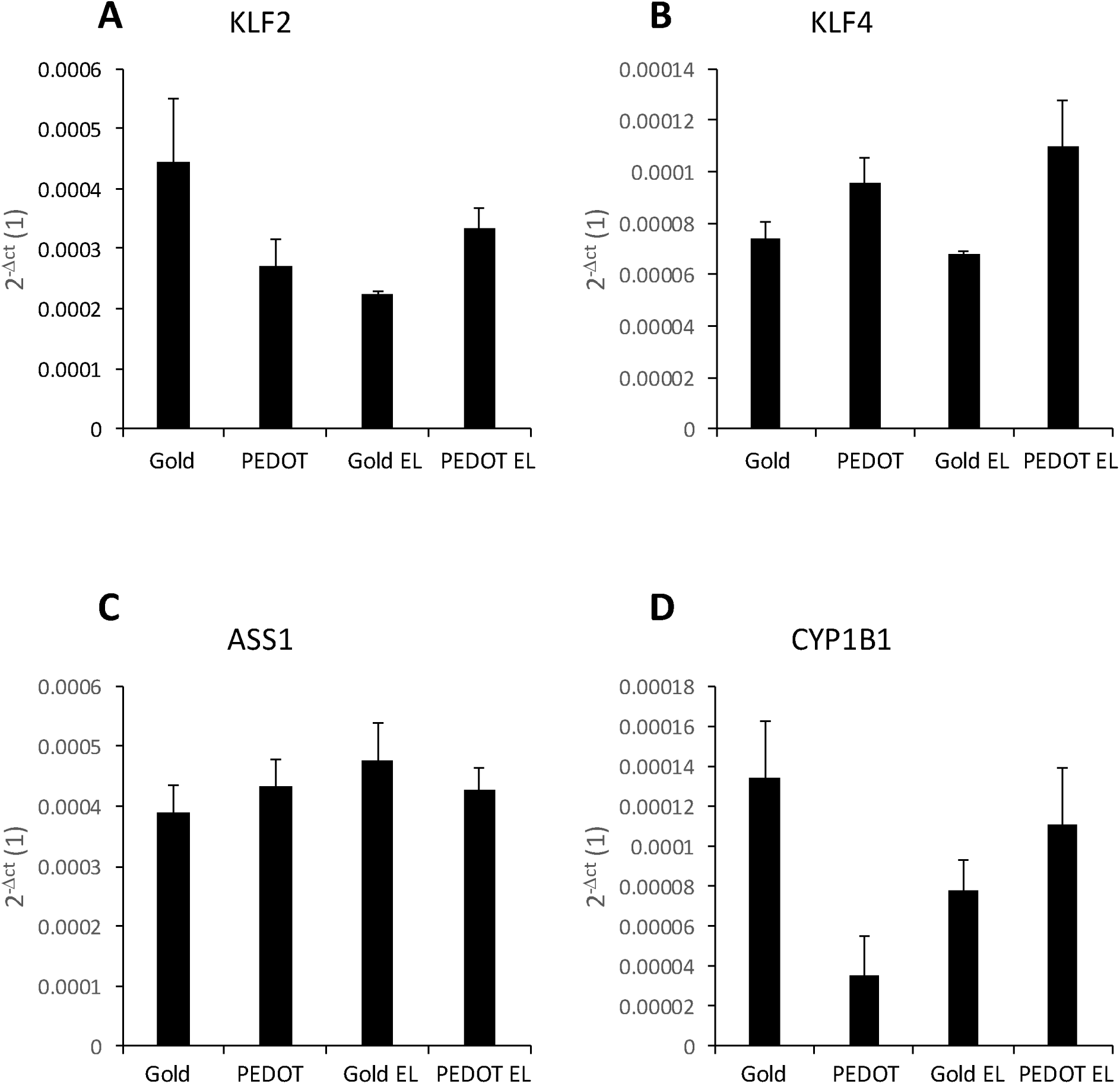
Expression of selected marker genes in cells. Human endothelial cells (HUVEC) were electostimulated for two days. mRNA transcripts of Krüppel-like factor 2 (KLF2, **A**), Krüppel-like factor 4 (KLF4, **B**), Argininosuccinate synthetase 1 (ASS1, **C)** and cytochrome 1B1 (CYP1B1, **D**) were determined by RT-qPCR. “EL” indicates electrostimulated variants. Data are indicated as mean value ± SEM, N=4-8. Significant differences (ANOVA, p < 0.05) are indicated.

Electrostimulation Effect: Electrostimulation tended to suppress KLF2 expression on gold electrodes, while having a negligible effect on PEDOT electrodes.

Material Effect: Without electrostimulation, KLF2 expression was lower on PEDOT electrodes compared to gold. However, with electrostimulation, the trend reversed, showing higher KLF2 expression on PEDOT electrodes than on gold.

#### KLF4 Expression (Fig. 7B)

Electrostimulation Effect: Electrostimulation had minimal effect on KLF4 expression for both gold and PEDOT electrodes.

Material Effect: PEDOT electrodes showed a trend of enhancing KLF4 expression compared to gold electrodes, regardless of electrostimulation.

#### ASS1 Expression (Fig. 7C)

No notable trends were observed in ASS1 expression across conditions, as both electrostimulation and electrode material had negligible effects.

#### CYP1B1 Expression (Fig. 7D)

Electrostimulation Effect: Electrostimulation appeared to suppress CYP1B1 expression on gold electrodes, while it tended to enhance CYP1B1 expression on PEDOT electrodes.

Material Effect: Without electrostimulation, CYP1B1 expression was lower on PEDOT electrodes compared to gold. However, with electrostimulation, CYP1B1 expression was higher on PEDOT electrodes, reversing the trend observed in the absence of stimulation.

## Discussion

The most significant result of this work is that pulsed electrostimulation, in combination with electrode material, can improve the physiological status of vascular endothelial cells grown under static conditions. PEDOT electrodes, in particular, demonstrated greater support for endothelial functionality. Specifically, electrostimulation enhanced NO production, as indicated by increased nitrite accumulation in the cell culture media. NO is a hallmark of healthy endothelial cells, mediating vasodilation critical to vascular homeostasis (10). While literature has previously shown the relationship between electrostimulation and vascular endothelial regeneration as reviewed by Chen et al. (4), to the best of our knowledge, this is the first study to suggest that electrostimulation can directly enhance NO production in endothelial cells.

Cells electrostimulated with both gold and PEDOT electrodes exhibited nitrite accumulation in the culture media, with PEDOT electrodes showing a trend toward being more supportive (Fig. 6). Interestingly, this trend in nitrite accumulation was not accompanied by a clear priming of endothelial cell metabolism toward NO production, as evidenced by the lack of significant changes in ASS1 transcript levels (Fig. 7A). ASS1 is known to be related to production of NO in endothelial cells under physiological conditions (12). This lack of ASS1 upregulation, despite increased nitrite accumulation, suggests alternative mechanisms, potentially involving post-transcriptional regulation.

Further analysis of additional markers of endothelial cell physiological status (KLF2, KLF4, CYP1B1, (11,12) revealed notable patterns in the combined effects of electrode material and electrostimulation (Fig. 7B-D). Electrostimulation generally enhanced the expression of these genes at the mRNA level on PEDOT electrodes but suppressed it on gold electrodes, with KLF4 being an exception, showing minimal sensitivity to electrostimulation. Material-specific effects further demonstrated that PEDOT was more supportive under electrostimulation, promoting endothelial functionality as inferred from trends in KLF2, KLF4, and CYP1B1 expression. Conversely, PEDOT tended to suppress gene expression compared to gold in the absence of stimulation, as reflected in KLF2 and CYP1B1 transcript levels. Similar differential effects of electrode material (PEDOT vs. gold) in combination with electrostimulation were observed in previous work (15). These results may suggest that the capacitive properties and surface characteristics of PEDOT electrodes create a more favorable environment for endothelial cell stimulation. Collectively, these data underscore the potential of PEDOT electrodes in vascular tissue engineering and regenerative medicine applications.

In terms of cell morphology, the electrostimulation of endothelial cells with gold or PEDOT electrodes under static conditions showed minimal effects. A slight trend toward alignment was observed under electrostimulation with PEDOT electrodes (Fig. 4), but it did not result in significant or consistent alignment of cells along the electrodes. Similarly, cell elongation, as assessed by the elongation index, revealed no substantial changes due to electrostimulation or electrode material (Fig. 5). These findings suggest that while endothelial cells may detect the effects of electrostimulation and electrode material, these stimuli were insufficient to drive notable morphological changes. This outcome contrasts with existing literature, where similar external electric fields induced clear alignment of endothelial cells along the electrodes (14,19). Such discrepancies could be attributed to differences in electrode design, field strength, or experimental conditions, highlighting the need for further detailed investigations.

The platform design provided a sufficient growth area for cells and included a 1 mm-wide observation window between electrodes (Fig. 1A), allowing for effective microscopic evaluation of cell morphology and behavior. This feature ensured precise assessment of the experimental outcomes.

Surface parameters may extensivelly impact cell culture eperiment. Hence, the electrode surface characteristics were characterised. The gold electrodes exhibited a smooth surface, whereas the PEDOT electrodes displayed submicrometer-scale roughness and a porosity of approximately 100 nm (Fig. 1BC), consistent with previous reports (20). This increased roughness and porosity significantly enhanced the electrode surface area, correlating with a markedly higher electrical capacitance in physiological media (∼1.4 mF for PEDOT vs. ∼10 µF for gold). The enhanced capacitance was reflected in the higher currents observed for PEDOT electrodes during cyclic voltammetry measurements. This capacitive behavior likely contributed to the favorable outcomes observed during electrostimulation, providing a more physiologically relevant stimulation pattern.

Cyclic voltammetry also revealed that PEDOT electrodes primarily exhibited capacitive behavior, enabling the application of 200 mV/mm pulses without significant faradaic reactions (see Fig. 2B). Importantly, no detectable hydrogen peroxide was produced during electrostimulation, a known cytotoxic by-product that can arise at higher voltages (18). The absence of hydrogen peroxide highlights the suitability of PEDOT electrodes for safe and effective electrostimulation under physiological conditions.

In summary, this study indicated the potential of PEDOT electrodes for promoting endothelial cell functionality through electrostimulation. The combined effects of electrostimulation and material properties suggest that PEDOT is a promising candidate for regenerative medicine, particularly in vascular tissue engineering.

### Study limitations

The major limitation of the presented study is that the study focuses on endothelial cells grown under static conditions. This does not fully replicate the dynamic physiological environment of blood flow, which influences endothelial behavior. Future studies should incorporate flow-based systems to better simulate *in vivo* conditions.

## Acknowledgements

The authors thank to Patricia Kittová, Petra Raptová and Michaela Pešková for technical assistance.

The research was supported by Czech Science Foundation Grant No. 21-01057S

The authors have declared that no competing interests exist.

